# Species ecology explains the various spatial components of genetic diversity in tropical reef fishes

**DOI:** 10.1101/2021.01.21.427590

**Authors:** Giulia Francesca Azzurra Donati, Niklaus Zemp, Stéphanie Manel, Maude Poirier, Thomas Claverie, Franck Ferraton, Théo Gaboriau, Rodney Govinden, Oskar Hagen, Shameel Ibrahim, David Mouillot, Julien Leblond, Pagu Julius, Laure Velez, Irthisham Zareer, Adam Ziyad, Fabien Leprieur, Camille Albouy, Loïc Pellissier

**Affiliations:** Landscape Ecology, Institute of Terrestrial Ecosystems, ETH Zürich, CH-8092 Zürich, Switzerland; Swiss Federal Institute for Forest, Snow and Landscape Research WSL, CH-8903 Birmensdorf, Switzerland; Genetic Diversity Centre (GDC), ETH Zürich, CH-8092 Zürich, Switzerland; CEFE, Université de Montpellier, CNRS, EPHE-PSL University, IRD, Univ Paul Valéry Montpellier 3, Montpellier, France; MARBEC, Université de Montpellier, CNRS, IFREMER, IRD, Montpellier, 34095, France; Centre Universitaire de formation et de recherche de Mayotte, Dembeni, 97660, France; Centre National de la Recherche Scientifique (CNRS), UMR 248 MARBEC, Montpellier, France; Department of Computational Biology, University of Lausanne, 1015 Lausanne, Switzerland; Swiss Institute of Bioinformatics, Quartier Sorge, 1015 Lausanne, Switzerland; Seychelles Fishing Authority, Mahe, Seychelles; Maldives Whale Shark Research Programme, Popeshead Court Offices, Peter Lane, York, Yorkshire, Y01 8SU, UK; Institut Universitaire de France, Paris, France; Wildlife Conservation Society, Madagascar Program, Antananarivo, Madagascar; Mafia Island Marine Park, Mafia, Tanzania; Ministry of Fisheries and Agriculture, Malé, Republic of Maldives; IFREMER, Unité Écologie et Modèles pour l’Halieutique, rue de l’Ile d’Yeu, BP21105, 44311 Nantes cedex 3, France

**Keywords:** coral reefs, ddRADseq, ecological trait, genetic diversity, tropical reef fishes, Western Indian Ocean

## Abstract

Intraspecific genetic diversity should be dependent on species ecology, but the influence of ecological traits on interspecific differences in genetic variation is yet to be explored. Generating sequenced data for 20 tropical reef fish species of the Western Indian Ocean, we investigate how species ecology influences genetic diversity patterns from local to regional scales. We distinguish between the *α, β* and *γ* components of genetic diversity, which we subsequently link to six ecological traits. In contrast to what is expected by the neutral theory of molecular evolution, we find that the *α* and *γ* components of genetic diversity are negatively associated with species abundance, which can be explained by larger variance in reproductive success in large populations and/or higher introgression in less frequent species. Pelagic larval duration, an important dispersal trait in marine fishes, is found to be negatively related to genetic *β* diversity, as expected by theory. We conclude that the neutral theory of molecular evolution may not be sufficient to explain genetic diversity in tropical reef fishes and that additional processes influence those relationships.

## BACKGROUND

Genetic diversity is central to many conservation challenges, such as species responses to environmental changes, ecosystem recovery, and the viability of endangered populations (e.g.[1]). Theory predicts that neutral genetic diversity is proportional to the mutation rate and the effective population size *Ne* [2], with higher genetic diversity occurring in populations with a larger effective size [3]. Assuming that *Ne* is proportional to census population size, this should translate into a positive relationship between neutral genetic diversity and species traits associated with census population size, such as body size and fecundity [4]. However, this expected relationship is not always clear in empirical data (e.g. [5,6]) and the effective to census size ratio (*Ne/N)* often departs from 1:1 under the influence of life-history and ecological traits [7]. For example, *Ne* was found to be smaller than census population sizes for marine fishes (e.g. [8]), suggesting that large population sizes alone may not be sufficient to explain genetic variation. This deviation, known as the Lewontin paradox [9] and more generally as the observed variation of genetic diversity among species, calls for more studies determining how species ecology shapes patterns of genetic variation across species.

Analogous to the taxonomic diversity of species assemblages [10], the intraspecific genetic diversity of groups of individuals can be expressed as three components related to different spatial scales: (*i*) *α* diversity, defined as the genetic diversity within a single local group of individuals; (*ii*) *β* diversity, defined as the genetic differentiation between local groups of individuals geographically separated, and consequently reflecting the degree of population genetic structure; and (*iii*) *γ* diversity, defined as the genetic diversity across all individuals of a defined region [11,12]. Previous studies highlighted the importance of life-history and ecological traits in shaping genetic diversity *sensu stricto* (i.e. the *α* and *γ* components as measured by heterozygosity, allelic richness and nucleotide diversity) and genetic structure (i.e. the *β* component as measured by genetic differentiation including fixation metrics, e.g. *FST*; [13]), but few integrated more than one spatial component to offer a general understanding of the processes shaping genetic diversity across scales (table S1).

The *α, β* and *γ* components of genetic diversity could be associated with different demographic processes (e.g. reproduction, longevity) that play out on different spatial scales [14,15]. In animals, local genetic *α* diversity has been linked to species longevity and reproduction rate (i.e. r or K strategies; [16]). Frequent reproduction causes mutations during gametogenesis and faster accumulation of variation within local groups of reproducing individuals [16]. Genetic *α* diversity has been also found to be correlated with population size (e.g. [17]), because larger population sizes may counter deleterious effects of inbreeding and should be, on average, more polymorphic [18]. Romiguier *et al*. [4] measured genetic *γ* diversity, which is related to parental investment and fecundity, from 10 individuals across the species range, and demonstrated a response similar to that for genetic *α* diversity. Genetic *β* diversity, between geographically distant groups, has been related to dispersal traits (e.g. [19–21]). For example, lower genetic differentiation is detected in more connected demes in freshwater fish [22]. The restricted movement of individuals limits gene flow and promotes dissimilarity in species allelic composition, especially for small populations or species with small effective population sizes [23]. However, to the best of our knowledge, patterns have never been investigated by considering the three components of genetic diversity for multiple species distributed across multiple geographical locations and spanning an ecological trait gradient.

Tropical reef fishes exhibit a wide array of ecological strategies [24] and inhabit relatively isolated reef patches, which makes them a good model system to investigate the link between ecological traits and the three components of genetic diversity. Specifically, tropical reef fishes differ in their reproductive rate and longevity, as well as their dispersal ability and geographical range size [25], which should translate into varying levels of *α, β* and *γ* genetic diversity across species. For example, in 13 species of damselfish, Gajdzik *et al*. [15] found the lowest levels of genetic *α* diversity in lower trophic levels. The link between dispersal and *β* genetic diversity in tropical reef fishes has been investigated in other studies, indicating an association with Pelagic Larval Duration (PLD: [26–28]) and reproductive strategy [21]. Less is known about how ecological traits might shape genetic *γ* diversity. Moreover, because demographic processes can shape more than one component of genetic diversity [29], investigations about how biological processes affect these metrics will provide new insight to their inter-relatedness.

Herein, we evaluated the relationship between a set of ecological traits and the three components of genetic diversity in tropical reef fishes using double-digest restriction-site-associated DNA sequencing (ddRADseq). Single Nucleotide Polymorphisms (SNPs) are useful for exploring spatial patterns of genetic diversity because they enable the detection of fine-scale spatial structure, which would not be detected using other traditional genetic markers (microsatellites, AFLP, allozymes; [30]). The use of RADseq is therefore particularly appropriate to study genetic diversity patterns in the marine realm because of the absence of strong physical boundaries.

We selected 20 tropical reef fish species, spanning a large variation in ecological traits and co-occurring in 4 regions of the Western Indian Ocean (WIO). We quantified the *α, β* and *γ* components of genetic diversity using the formalized framework of Jost [11], which we linked with important ecological traits for tropical reef fishes. We generated a set of expectations on the link between biological processes, species trait proxies and the genetic diversity component under the Hardy-Weinberg equilibrium (table 1). We formed the following expectations: (1) Genetic *α* diversity is associated with species traits related to reproduction and population size. High reproductive outputs lead to larger population sizes, which favour mutation, reduce inbreeding probability and the effect of random drift, and hence increase local genomic diversity. (2) Genetic *β* diversity is associated with ecological traits related to dispersal ability that modulate gene flow, such as PLD influencing larval gene flow and adult body size affecting adult gene flow. (3) Genetic *γ* diversity is shaped by the circulation of genetic variation that arises locally across the species range and is associated with traits related to reproduction, dispersal and population size. As a corollary, deviation from these expectations would point to alternative processes possibly influencing genetic diversity in tropical reef species. By exploring different spatial components of genetic diversity from a large number of SNPs over multiple tropical reef fishes, our main objective was to shed light on the role of ecology in shaping genetic variation across species.

**table 1.** Expected relationships between the *α, β* and *γ* spatial components of genetic diversity and demographic parameters under Hardy Weinberg Equilibrium.

| Demographic process | Biological trait | Genetic process | $\alpha$ | $\beta$ | $\gamma$ |
| --- | --- | --- | --- | --- | --- |
| Population size | Census size/abundance | large population size: decrease in **genetic drift** | Increase<sup>[81]</sup> |  | Increase<sup>[33]</sup> |
|  | Adult body size | small species have larger population sizes: decrease in **genetic drift** | Decrease<sup>[5]</sup> |  | Decrease<sup>[82]</sup> |
|  | Schooling | large group size: decrease in **genetic drift** | Increase |  | Increase |
| Reproduction | Parental investment/reproduction | large fecundity (lower investment): increase in population size and decrease in **genetic drift** | Increase | Decrease<sup>[83]</sup> | Increase<sup>[83]</sup> |
|  | Adult body size | larger species with lower reproduction: decrease in **mutation rates** | Decrease |  | Decrease<sup>[4,84]</sup> |
| Dispersal | Pelagic larval duration | long larval duration: increase in **gene flow** |  | Decrease<sup>[84]</sup> | Increase<sup>[85]</sup> |
|  | Home range | large species home range: high **gene flow** |  | Decrease | Increase<sup>[86]</sup> |
|  | Adult body size | large species body size: **high gene flow** |  | Decrease | Increase |

## MATERIAL AND METHODS

### Species selection along gradients of ecological traits

We selected 20 tropical reef fish species from the WIO, ranging from small- to large-bodied, short to long PLDs and including various abundances on the reef (see table S2), providing a valid representation of the ecological variation of the entire trait space [31]. We ran a Principal Component Analysis (PCA) over all species co-occurring in the target sampling locations (n=2292) using five ecological traits gathered from Fishbase [32] and the literature (see Supplementary methods), namely: (i) adult body size (cm); (ii) PLD (days); (iii) adult home range mobility; (iv) reproductive guild (guarders vs. non-guarders); and (v) schooling. Then, for these 20 selected species, we considered an estimation of their regional-scale species abundance as a proxy of population census size (see [33]), i.e. the cumulated number of individuals per transects over the WIO, gathered from the Reef Life Survey (RLS) program (https://reeflifesurvey.com). We tested the correlations among all the traits (figure S2) and investigated the phylogenetic signal of these traits by using the λ metric [34] for continuous traits and the D-statistic [35] for binary traits (table S3).

### Field sampling

In 2016–2017, we sampled a total of 852 individuals across the 20 target species from different sites in four sampling locations (figure 1), in the Republic of Maldives, Tanzania (Mafia Island), the Seychelles and the Comoros Archipelago (Mayotte Island). We performed the sampling by SCUBA diving using hand barrier nets. For the largest-bodied and most mobile target species, we complemented the sampling with fish from local fish markets. For each specimen, muscle tissue was collected and stored in either 90% EtOH or RNAlater until processing.

**figure 1.**
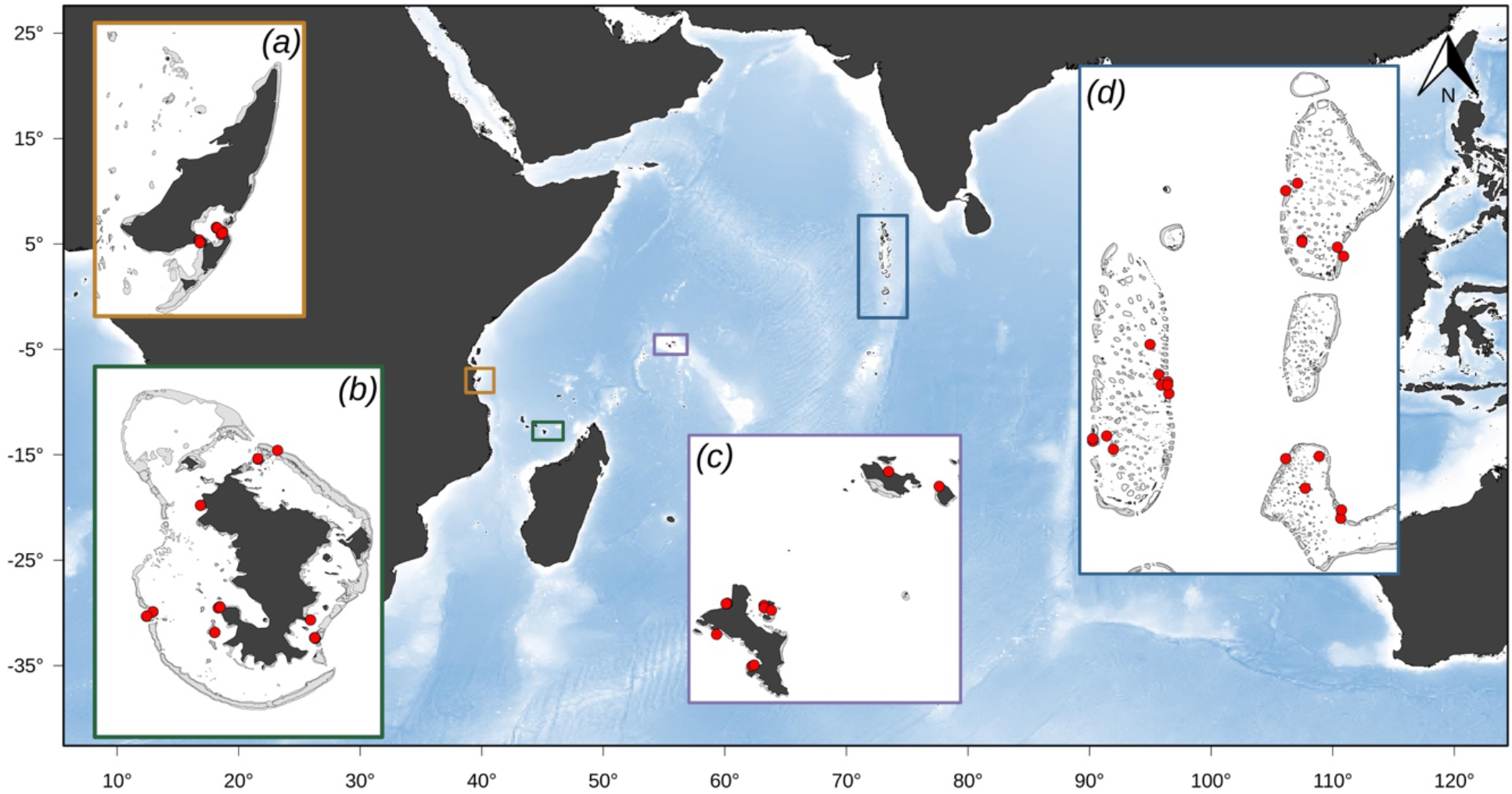
Map of the sampling locations in the Western Indian Ocean: Mafia Island (Tanzania; *a*), Mayotte Island (Comoros, France; *b*), the islands of Mahe, Praslin and La Digue (Seychelles; *c*) and three central atolls of the Maldives, namely Kaafu North, Alif Alif, Vaavu and Kaafu South (Republic of Maldives; *d*). Specific sampling sites are indicated by red points.

### Genotyping

High-quality genomic DNA was extracted from muscle tissue using the LGC, sbeadex livestock kit (catalog numbers 44701 and 44702). ddRAD-seq libraries were prepared using EcoRI and Taq1a (New England Biolabs, Inc., Ipswich, MA, USA) following the protocol used in Westergaard *et al*. [36]. 24 ddRAD-seq library pools containing 2 x-48 internal barcodes each were sequenced in 12 lanes on the Hiseq 2500 Illumina platform using the 2 × 125 bp protocol (Fasteris, Geneva, Switzerland). We used the default settings of the dDocent pipeline v.2.2.25 [37] to obtain the genotypes. Raw reads were demultiplexed using Stacks (v.2.0b: [38]). Per species, we built a reference catalog *de novo*. In order to find the optimal parameters, we maximized the remapping rate by varying the coverage of unique sequences within individuals, the number of shared loci among samples, and sequence identity (table S4). Reads were remapped to the reference catalogs using BWA v.0.7.17 [39] and SNPs were called using FreeBayes v.1.3[40]. Total SNPs were filtered using VCFtools v.0.1.16 and vcflib v.1.0.1[41]. We only kept SNPs that had been successfully genotyped with a minimum quality score of 20, minimum mean depth of 3, mean depth of 10, minor allele count of 3, and minor allele frequency of 5%. We removed loci with more than 20% missing data per population. We filtered for allele balance and mapping quality between the two alleles. We removed loci with coverage that was too high, decomposed complex SNPs into single SNPs, removed indels and sites with missing data (> 5%), and kept only biallelic sites in Hardy-Weinberg equilibrium. Finally, we used RAD haplotyper [42] with the default settings to remove putative paralogous loci. A Principal Component Analysis (PCA) was conducted on the SNP data for each species using the *glPca* function implemented in the “adegenet” package [43] in R v.3.6.0 (R Core Team 2019; figure S3), overall, we identified 50 outliers (mean = 1.92 per species) across the sequenced individuals and removed them prior downstream analyses. For the comparison of the 20 investigated species, we accounted for differences in population sampling success by standardizing the sample size to a maximum number of 10 individuals per population (median and tradeoff value of the overall sampling). We also down-sampled the filtered SNP data 999 times to the lowest common number of SNPs (i.e. 4479) found across all species (table S5), a necessary procedure given the link between SNP number and genetic diversity[44].

### Genetic 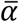, ***β* and *γ* diversity**

To quantify the different components of genetic diversity (*α, β* and *γ*), we applied the multiplicative partitioning framework proposed by Jost [11] for genetic diversity, expressed as follows: *J_T_* = *J_S_* × *J_ST_*. Three equations within this framework are based on Hill’s number, and enable the partitioning for true diversities, as done in Gaggiotti *et al*. [12]. In this framework, *J_ST_* represents the mean within-population genetic component 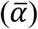 and can be expressed as the expected mean heterozygosity of populations (*H_s_*: Nei [45]) as follows: 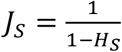. *J_ST_* represents the between-population (*β*) component and can be expressed as 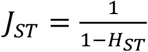 where 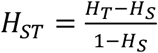 and *H_T_* represents the overall genetic diversity. We calculated *H_T_* and *H_s_* corrected by the number of individuals for each species and each of the 999 × 4479 SNP data sets using the *basic*.*stats* function implemented in the R package “hierfstat” [46]. We compared the multiplicative framework to an additive one, where 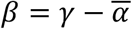. Similarly, for this framework, we used the mean heterozygosity (*H*_s_) as measure of 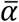 diversity and the overall gene diversity (*H*_T_) as a measure of *γ* diversity. The *β* diversity component is equivalent to the *D*_ST_ metric, where *D_ST_* = *H_T_* – *H_s_* [45]. All these metrics were calculated for each species and each of the 999 × 4479 SNP data sets. The quantification of the additive framework can be found in table S6. It shows that the metrics derived from the multiplicative and additive frameworks are strongly correlated (i.e. Pearson’s correlation coefficient: *rp* > 0.99; figure S2). Several metrics have been proposed to estimate the level of genetic differentiation among populations, namely *D_ST_’*[45], *F_ST_* [47], *F_ST_’, G_ST_* [45], *G_ST_’, G_ST_’’*[48] and *Dest* [11].We therefore compared these metrics to those derived from the multiplicative (*J_ST_*) and additive (*D_ST_*) partitioning approaches and found strong correlations (Pearson’s correlation coefficient: *rp* > 0.99; figure S2).

Based on these metrics, we further investigated the spatial variation in 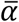 genetic diversity across the four sites and across species (figure S4*a*; table S7), as well as spatial patterns of *β* genetic diversity across sites (table S8) and in relation to geographical distance (figure S4*b*). To do so, we calculated the pairwise genetic *β* diversity (*F_ST_*) using the function *genet*.*dist* implemented in the R package “hierfstat” [46]. To explore the relationship between pairwise genetic *β* diversity and geographical distance, we fitted a Generalized Linear Model (GLM) with a negative exponential function, which describes the increase in species dissimilarity with increasing spatial distance [49] using the *decay*.*model* function in the R package “betapart” [50].

### Relationships between genetic diversities and ecological traits

We assessed the relationships between the 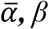 and *γ* components of genetic diversity and each ecological trait, using Phylogenetic Generalized Least Squares (PGLS) models to account for the phylogenetic non-independence of species [51]. The use of Ordinary Least Squares (OLS) models in phylogenetic comparative analysis can lead to type I error when assessing the significance of the regression coefficients. A regression model assuming a Gaussian error is justified here since the genetic diversity metrics are derived from Hill’s number, which follows a normal distribution. We used the *pgls* function implemented in the R package “caper” [52] and the time-calibrated phylogeny of [53]. The *pgls* function enables the estimation (via maximum likelihood) of a phylogenetic scaling parameter, which indicates the degree of phylogenetic dependency in correlations among the response and explanatory variables. For data sets with few sampled specimens, the estimation of Pagel’s λ by maximum likelihood does not perform well [51]. We therefore report the results of PGLS models with λ = 1 (complete phylogenetic dependence) but also those with λ = 0 (phylogenetic independence), which corresponds to OLS models. For quantitative traits, we report the regression coefficient together with its *P*-value, while for qualitative traits we report the Fisher statistic (F) derived from an ANOVA applied to the PGLS model, along with its *P*-value. Last, for each genetic diversity component, we extracted the corrected Akaike’s information criterion (AICc) from the six PGLS models to compare their relative goodness of fit, where the model with the lowest AICc is considered the estimated best model. From the set of AICc values, we calculated the Akaike weight of each PGLS model [54]. The Akaike weight, ranging from 0 to 1, is interpreted in terms of the probability of a given model being the best in explaining the data within a predefined set of alternative models. This metric made it possible to identify which of the six considered traits best fit each genetic diversity component. Given the small number of observations (n = 20) and small number of explanatory variables (n = 6), which can lead to overfitting and spurious relationships, we did not consider a full PGLS model including all traits together. We used a logarithmic transformation for abundance and adult body size, as preliminary analyses showed that it significantly improved the linearity of the relationships.

Because the six studied traits are not independent (see figure S1), translating into different trait syndromes, we also performed the OLS and PGLS models using as explanatory variables the coordinates of the 20 species on the first (PC1) and second (PC2) axes of a Principal Component Analysis, accounting for circa 80% of the trait variation among species (figure S3). PC1 (explaining 65.1% of the variance) reflects a gradient of abundance on the reef, body size, reproductive guild, home range mobility behaviour and PLD, with positive coordinates including species with lower abundances, larger body size, higher PLD, wider home range mobility and adopting a “non-guarder” reproductive strategy. While, PC2 (explaining 14.6% of the variances) only associated with schooling, includes species gathering in larger schools at highest PC2 values.

## RESULTS

### Genome-wide SNP data for 20 tropical reef fish species

We genotyped 20 tropical reef fish species across four sampling locations for a total of 852 individuals at 14,740 ± 7633 SNPs across the genome (table S5). Considering the multiplicative framework, we found no significant relationship between 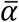 and *β* genetic diversity (PGLS: R^2^ = 0.016, coefficient = 0.97, *P* = 0.59), nor between *β* and *γ* diversity (PGLS: R^2^ = 0.091, coefficient = 2.38, *P* = 0.20). We found a very strong positive association between 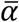 and *γ* genetic diversity (PGLS: R^2^ = 0.97, coefficient = 0.95, *P* < 0.001). Altogether, the three different genetic components can be summarized in a triangular space (figure 2), covering the species’ overall genetic diversity properties along with the most important species ecological traits (see below). Because 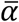 and *γ* genetic diversity are highly correlated, this representation allows the identification of three main groups. The first group included species with a small adult body size, short PLD and medium abundance, presenting low values of 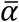 and *γ* diversity but medium values of *β* diversity, with species such as *Chromis weberi, Chromis atripectoralis* and *Dascyllus trimaculatus* as representatives (figure 2). The second group encompassed species with a medium body size, short PLD and low abundance, presenting medium values of 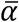 and *γ* diversity but low values of *β* diversity. In this group we found species such as *Myripristis violacea, Ctenochaetus striatus* and *Gomphosus caeruleus* (figure 2). The third group, represented by species such as *Caranx melampygus* and *Naso brevirostris* (figure 2), represents species with a large body size, long PLD and low abundance, which were associated with high values of 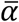 and *γ* diversity and very low values of *β* diversity.

**figure 2.**
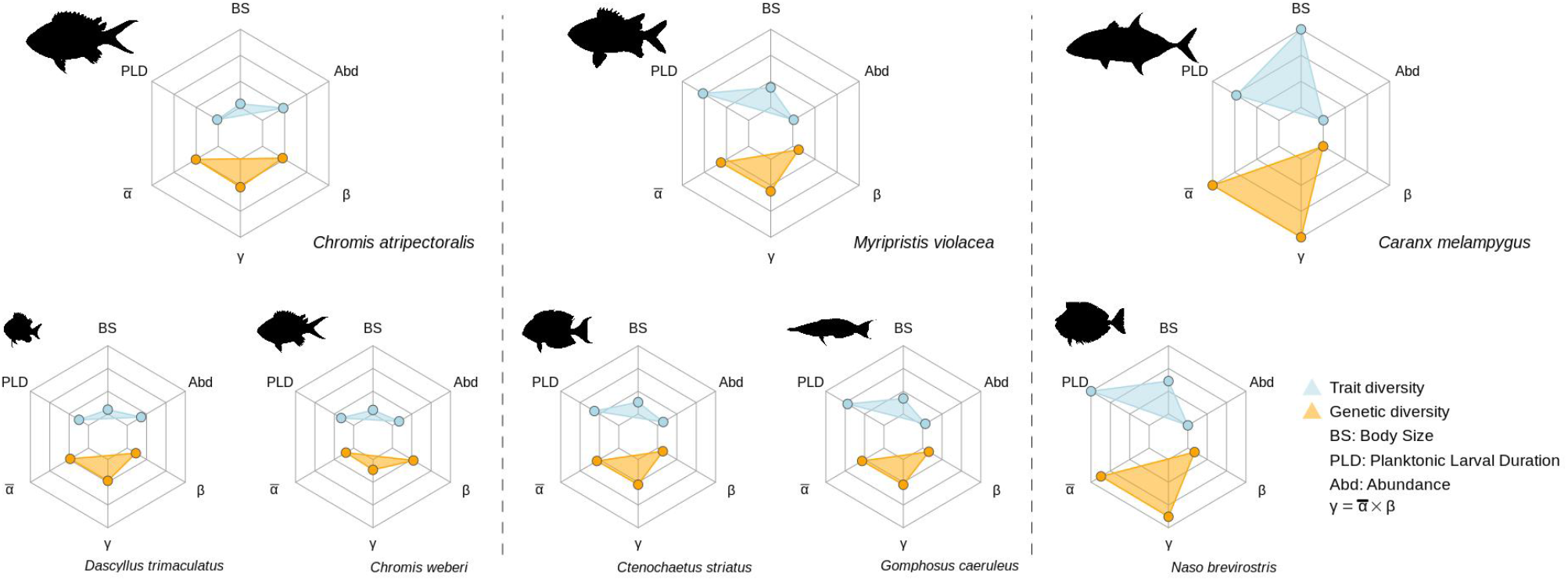
Radar plots illustrating the components of genetic diversity for selected tropical reef fishes with different ecological traits. The first group is composed of species with a small body size (BS), short pelagic larval duration (PLD) and medium abundance (Abd), presenting low values of 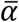 and *γ* diversity but medium values of *β* diversity, such as *Chromis weberi, Chromis atripectoralis* and *Dascyllus trimaculatus* (all three Pomacentridae). The second group encompasses species such as *Myripristis violacea* (Holocentridae), *Ctenochaetus striatus* (Acanthuridae) and *Gomphosus caeruleus* (Labridae), with a short PLD, medium body size and low abundance, which present medium values of 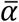 and *γ* diversity but low values of *β* diversity. The third group comprises species with a low abundance, long PLD and large body size, which are associated with high values of 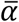 and *γ* diversity and very low values of *β* diversity. This group is characterized by species such as *Caranx melampygus* (Carangidae) and *Naso brevirostris* (Acanthuridae).

### Genetic *α* diversity and ecological traits

No significant differences in genetic *α* diversity were found between sites (Kruskal-Wallis chi-squared = 3.580, DF = 3, *P* = 0.311; figure S4*a*). When considering the 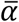 genetic diversity across the four sites, the PGLS model with species abundance as a predictor provided the best support (W_AICc_ = 0.63; table 2, figure 3) among the six alternative models (table 2). 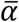 genetic diversity was negatively associated with species abundance (PGLS: R^2^ = 0.479, coefficient = -0.0138, *P* = 0.0009; figure 4, table 2) and positively associated with adult body size (PGLS: R^2^ = 0.431, coefficient = 0.0380, *P* = 0.002). More precisely, *Caranx melampygus* and *Naso brevirostris* showed the highest values of *J_S_*, with 1.474 and 1.446, respectively, while the lowest values were found for *Pseudanthias squamipinnis* and *Chromis weberi*, with 1.309 and 1.332, respectively (table S6). Less abundant species on the reef, with a large body size, such as the solitary *Hemigymnus fasciatus* (*J_S_* = 1.424; table S2 for ecological traits) and *Caranx melampygus* (*J_S_* = 1.472; table S6) had higher *J_S_* values in comparison to more abundant species with a smaller body size, such as *Dascyllus carneus* (*J_S_*= 1.347) or *P. squamipinnis* (*J_S_* = 1.313, table S6).

**table 2.** Full results of the phylogenetic generalized least squares (PGLS) models for the 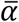, *β* and *γ* components of genetic diversity and the six ecological traits investigated. For quantitative traits, we report the significance of the PGLS regression coefficient (slope), while for qualitative traits we report the significance of the Fisher statistic (F) derived from an ANOVA applied to the PGLS model. Significant variables (*P* < 0.05) are given in bold.

| $\bar{\alpha}$ genetic diversity | PGLS | | | | | |
| --- | --- | --- | --- | --- | --- | --- |
|  | R<sup>2</sup> | Coef. | F | P | AICc | W<sub>AICc</sub> |
| **Body size** | **0.431** | **0.0380** | - | **0.00167** | **-84.850** | **0.321** |
| Pelagic larval duration | 0.0474 | 0.00051 | - | 0.356 | -74.552 | 0.002 |
| **Abundance** | **0.468** | **-0.0138** | - | **0.00088** | **-86.197** | **0.629** |
| Home range | 0.307 | - | 7.961 | 0.011 | -80.905 | 0.045 |
| Schooling | 0.0332 | - | 0.618 | 0.442 | -74.255 | 0.002 |
| Reproduction | 0.0451 | - | 0.850 | 0.369 | -74.503 | 0.002 |
| $\beta$ genetic diversity | PGLS | | | | | |
|  | R<sup>2</sup> | Coef. | F | P | AICc | W<sub>AICc</sub> |
| Body size | 0.0265 | -0.00124 | - | 0.493 | -155.06 | 0.066 |
| **Pelagic larval duration** | **0.199** | **-0.00124** | - | **0.0483** | **-158.97** | **0.468** |
| Abundance | 0.147 | -0.00124 | - | 0.0952 | -157.70 | 0.248 |
| Home range | 0.0199 | - | 0.365 | 0.553 | -154.92 | 0.062 |
| Schooling | 0.00370 | - | 0.0663 | 0.800 | -154.60 | 0.052 |
| Reproduction | 0.0700 | - | 1.355 | 0.260 | -155.97 | 0.104 |
| $\gamma$ genetic diversity | PGLS | | | | | |
|  | R<sup>2</sup> | Coef. | F | P | AICc | W<sub>AICc</sub> |
| **Body size** | **0.362** | **0.0363** | - | **0.005** | **-80.90** | **0.049** |
| Pelagic larval duration | 0.0003 | 0.0003 | - | 0.587 | -72.25 | 0.001 |
| **Abundance** | **0.525** | **-0.0152** | - | **0.0003** | **-86.80** | **0.938** |
| **Home range** | **0.258** | - | **6.242** | **0.0223** | **-77.87** | **0.011** |
| Schooling | 0.0272 | - | 0.503 | 0.487 | -72.46 | 0.001 |
| Reproduction | 0.0249 | - | 0.460 | 0.506 | -72.42 | 0.001 |

**figure 3.**
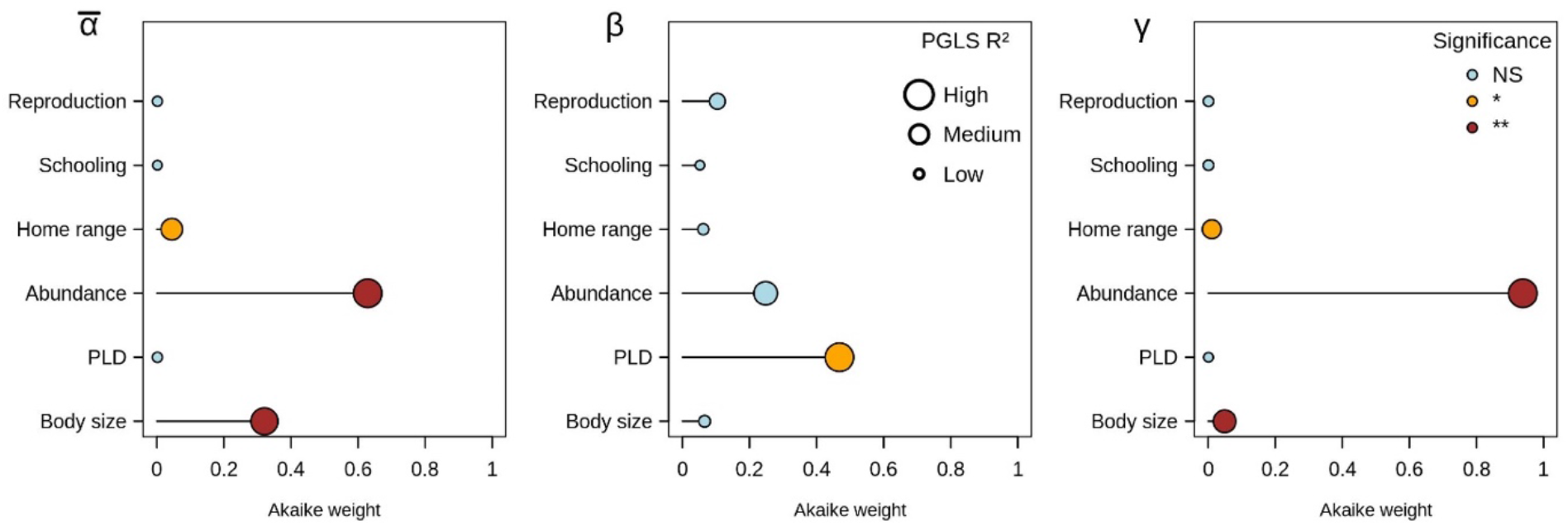
Relative support of each phylogenetic generalized least squares (PGLS) model in explaining the variation in the 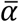, *β* and *γ* components of genetic diversity using a multiplicative framework of diversity partitioning. A PGLS model was fitted for each species trait separately and the corresponding Akaike weight, ranging from 0 (minimum support) to 1 (maximum support), was extracted. For quantitative traits we report the significance of the PGLS regression coefficient (slope), while for qualitative traits we report the significance of the Fisher statistic (F) derived from an ANOVA applied to the PGLS model, with ** for *P* < 0.01 and * for *P* < 0.05 (see the main text and table 2 for full results). The size of each circle is proportional to the R^2^ extracted from the PGLS model.

**figure 4.**
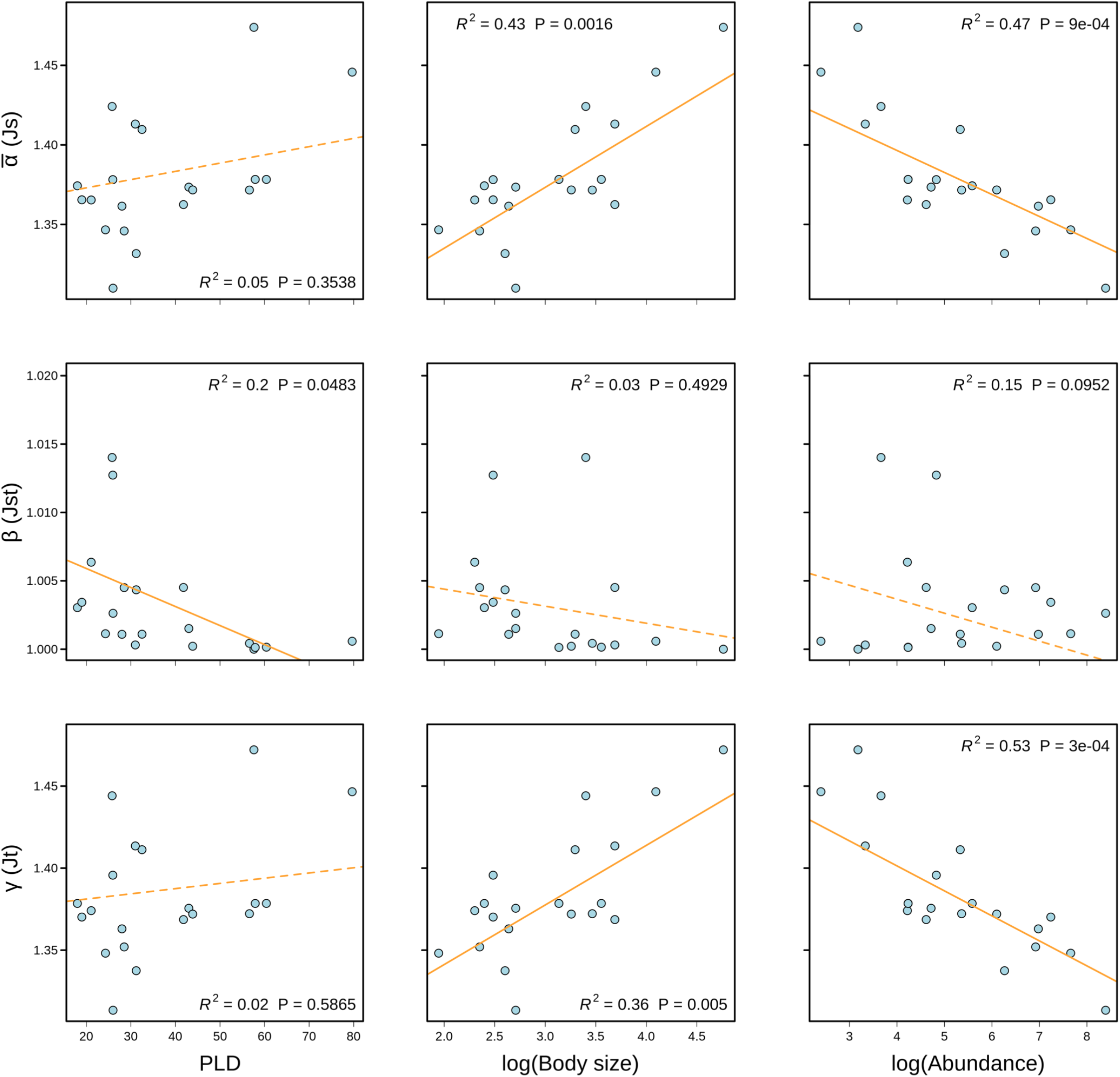
Relationships between the 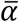, *β* and *γ* components of genetic diversity considering the multiplicative framework and the traits that provide the greatest support in phylogenetic generalized least squares (PGLS) models according to the Akaike weight (pelagic larval duration [PLD], adult body size and abundance). The orange lines represent the slopes estimated by PGLS models. A solid line indicates a significant relationship, while a dotted line indicates a non-significant relationship.

### Genetic *γ* diversity and ecological traits

Genetic *γ* diversity showed a similar association with ecological traits than that observed when considering genetic 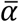 diversity. The PGLS model with abundance as a predictor provided the best support (W_AICc_ = 0.94; table 2, figure 3) among the six alternative models for genetic *γ* diversity (figure 4, table 2). Genetic *γ* diversity and species abundance had a significant negative relationship (PGLS: R^2^ = 0.525, coefficient = -0.0152, *P* = 0.0003; figure 4, table 2). Moreover, genetic *γ* diversity displayed a significant positive association with adult body size (PGLS: R^2^ = 0.362, coefficient = 0.0366, *P* = 0.005; figure 4, table 2). In particular, the lowest values were found for small-bodied species, such as *P. squamipinnis* and *Chromis weberi* (*J_S_* = 1.337), while the highest values were found for large-bodied species, such as *Caranx melampygus* and *N. brevirostris*, reaching, *J_S_* = 1.472 and 1.447, respectively (table S6). Species with a low home range mobility displayed lower levels of genetic *γ* diversity than those with a large home range mobility (PGLS: F = 6.242, *P* = 0.0223; table 2, table s9).

### Genetic *β* diversity and ecological traits

For most of the studied species, we found an association between pairwise values of genetic *β* diversity and the geographical distance between sites (figure S4*b*). Indeed, species with a shorter PLD, such as *Hemigymnus fasciatus* (PLD = 25.8; GLM: slope = 2.413e-05, *P* = 0.01) and *Oxymonacanthus longirostris* (PLD = 25.95; GLM: slope = 1.611e-05, *P* = 0.01), present higher levels of genetic differentiation in relation to geographical distance than species with a longer PLD, such as *Caranx melampygus* (PLD = 57.6; GLM: slope ≈ 0, *P =* 0.44) and *Zanclus cornutus* (PLD = 57.9; GLM: slope = 1.656e-07, *P* = 0.11; figure S4*b*). Results from the comparison of the PCA run on all the SNP data indicated that geographical differentiation among individuals ranged from weak, in five species (e.g. *Z. cornutus*; figure S5), to comparatively strong in *O. longirostris* (figure S5). For 13 species, we observed a gradient of genetic differentiation along the first component of the PCA (figure S3), reflecting a gradual and species-specific isolation between individuals in different geographical locations (figure S5). This gradient systematically isolated individuals from the Maldives from individuals coming from other sampling locations (figure S5). The PGLS model with PLD as a predictor provided the best support (W_AICc_ = 0.47) among the six alternative models for genetic *β* diversity (figure 3, table 2). Genetic *β* diversity displayed a significant negative association with PLD (PGLS: R^2^ = 0.200, coefficient = -0.00014, *P* = 0.048; figure 4; table 2). Species with the shortest PLDs (< 4 weeks), such as *Canthigaster valentini* and *O. longirostris*, were more differentiated than species with PLDs more than twice as long, such as *M. violacea* and *Z. cornutus*, which had highly connected populations (figure S3, table S8, S9).

### Genetic diversity and trait syndromes

We investigated the correlations between species traits, and their association into distinct trait syndromes (see figure S3, i.e., large bodied species with low regional abundance vs. small bodied species with high regional abundance) as well as their association with genetic diversity. Both PGLS models revealed a significant negative association between the 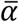, and γ components of genetic diversity and the PCA axis 1 (table S10), which confirms that large-bodied species with low abundance display the highest levels of 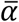 and *γ* genetic diversity.

## DISCUSSION

By considering the multiple components of genetic diversity, in analogy to the concept of species diversity in community ecology [10], our study shows that genetic diversity *sensu stricto* (*α* and *γ* components) and genetic differentiation (*β* component) are associated with different ecological traits acting at different spatial scales. We demonstrate that ecological traits can partly explain differences in genetic diversity among species, even within a restricted taxonomic range of species (e.g. [6,21,55]). In contrast to our expectations, both the *α* and *γ* components of genetic diversity were found to be negatively related to species abundance, a proxy for population size [17], and positively related to adult body size, which is considered an integrative biological trait in fishes [56]. In contrast, *β* diversity was found to be only associated with PLD, which is one of the main dispersal traits in tropical reef fishes [25]. Our results suggest that considering simultaneously multiple spatial components of genetic diversity leads to a better understanding of the processes shaping genetic diversity within and across species.

Larger effective population sizes and presumably larger local species abundances on reefs are amongst the factors expected to be positively associated with genetic diversity [3,17]. In contrast to what is expected by the neutral theory and suggested in previous studies (e.g.[17,33]), we found that both *α* and *γ* components of genetic diversity were negatively related to regional-scale abundance estimated from pooled underwater visual surveys over the West Indian Ocean. Factors that can lead to significant deviations from neutral theory in finite populations reviewed previously [57]. Temporal fluctuations in population size, as well as variation in reproductive success among individuals, can to a certain extent explain the discrepancy between our observations and theory [7]. Large variation in individual reproductive success in very large groups may translate into differences in allelic frequencies and generate a lower genetic diversity [58]. Further, marine species with high fecundity and high early mortality may also display high variance in reproductive success among individuals as a consequence of stochastic factors, which makes successful reproduction a ‘sweepstakes’ (e.g. Christie *et al*.[59] for tropical reef fishes). In support of this concept, we showed that the sea goldie (*P. squamipinnis*), known to have ‘sweepstakes’ reproduction and to form large aggregations [60], displayed the highest abundance but also the lowest levels of both *α* and *γ* genetic diversity. Moreover, Richards & van Oppen [61] found that rarer species can have very high genetic diversity in corals, suggesting that hybridization could explain higher than expected levels of genetic diversity in less frequent species. Hybridization also occurs in tropical fishes, and is associated with external fertilization [62] and competition for limited spawning grounds [63], but rarity is also expected to be an important factor (e.g. Frisch & van Herwerden [64]). Lower species abundance on the reef could favour interbreeding, because of the lower likelihood of finding conspecific partners [65], and as a result could increase genetic variation within those species. Despite earlier evidence of a positive relationship between local abundance and genetic diversity for pelagic fishes [66], our results suggest that expectations from theoretical models [2] might not be readily transferable to more complex and dynamic reef fish systems, where multiple biological processes can interact.

We also found that both the *α* and *γ* components of genetic diversity were positively related to adult body size, a trait typically considered as a proxy for population size [6], which was found to be inversely related to regional-scale species abundance in this study. Our result contrasts with expectations from the neutral theory and from the previously documented negative relationship found in studies targeting broader taxonomic scales [4]. In tropical reef fishes, adult body size integrates many ecological aspects, other than population size, such as habitat use and reproductive strategy [25], trophic level [67], predation [68], and ecological generality [25]. Furthermore, large-bodied species generally have a longer life span and a longer active adult dispersal period, also implying a larger range size [25]. These traits might make them able to pool genetic variations arising from different locations, thereby increasing overall genetic *α* and *γ* diversity and decreasing the effect of local drift that could cause a range-wide decline in genetic diversity.

The negative association between genetic *β* diversity and PLD detected in our analyses was expected, particularly for benthic marine species with predominantly immobile adults. For those species, dispersal is mostly the result of larval duration, which makes a considerable contribution to gene flow between geographically isolated populations [20]. Similar relationships were detected in previous studies on single species (e.g. [69,70]), but also in a multi-species context (e.g. Selkoe *et al*. [27]for tropical reef fishes). Meta-analyses, however, have given mixed results (e.g. [71]). The varying effect size among studies might be due to a scale dependence of the influence of dispersal [5] and to extrinsic factors such as geographical distance, seascape heterogeneity and barriers to dispersal related to ocean circulation and geological history (e.g. [72]). Better surrogate than PLD might be quantified in the future and will probably show more explanatory relationships. In particular, a meta-analysis showed that larval swimming capacities measured as mean critical swimming speed (U-crit) best explained genetic differentiation and range size in demersal fishes [73]. Overall, our results emphasize that, in the Western Indian Ocean, larval dispersal partly explains genetic differentiation between groups separated by large geographical distances (figure. S4b). However, more studies are needed to shed light on how dispersal processes and seascape features interact in shaping genetic differentiation in tropical reef fishes.

While our results provide novel insight on the link between genetic diversity and species ecology, our study has limitations relating to the number of sampling locations and design. First, *α* genetic diversity was estimated assuming that sampling groups were the unit of the analysis. Similar strategies were adopted by Wang *et al*. [74] for lizards and Dapporto *et al*. [75] for butterflies to detect drivers of *β* diversity among species. While using a genetic pool provides the best solution to define *α* genetic diversity [76], this approach is not possible when comparing multiple species with distinct genetic spatial structures. Consequently, for species with low genetic structure, *α* diversity will resemble *γ* diversity. Second, the application of nuclear SNP markers, which are still underused in research [15], makes comparisons with other studies based on mtDNA or microsatellite markers difficult. We conclude that the use of SNPs in a spatial context is promising, helps to detect more subtle genetic structures, and shows more subtle associations with ecological traits [77]. Finally, our study was limited in terms of the number of sampled sites, but we included a large set of species belonging to different clades and spanning a large ecological diversity gradient. Hence, our design is robust to investigate the link between species genetic components and ecological traits, but it could be expanded to evaluate whether the trait–genetic diversity relationships that we observed are still consistent across more sites in the WIO, and more generally throughout the Indo-Pacific region.

We linked the genetic diversity and ecology of tropical reef fishes, shedding light on the role of demographic and dispersal processes in shaping spatial patterns of genetic diversity. Small bodied species that are more abundant regionally, typically “benthic guarders” and with a low home range mobility, showed the lowest levels of *α* and γ genetic diversity. Because of lower genetic diversity, these species might be the least able to adapt under environmental changes such as climate change and over-harvesting [78]. However, their large population sizes and the maintenance of gene flow may help to maintain their adaptive potential by enhancing the overall genetic diversity [78]. Dispersal between reef patches is critical for the replenishment of individual populations, and our results support the use of PLD as a proxy for genetic connectivity among populations at the regional scale, and its use for conservation schemes [79]. Finally, our findings suggest that the degree of intraspecific genetic diversity may result not only from neutral demographic processes associated with population size [2], but also from additional ecological processes associated with species natural history. Beyond considering intrinsic ecological factors, future studies could investigate how ecological traits of species interact with external factors, including seascapes and historical dynamics, to shape levels of genetic diversity in species, and focus on the adaptive component of genetic diversity [80].

## ETHICS

The collected data has no commercial value and cannot be used in a way that could be detrimental to local populations. Sampling was performed in accordance with local regulations and with local collaborators, research permits number are: Maldives (OTHR) 30-D/INDIV/2016/538), Mayotte (06/UTM/2016), Seychelles (A0157) and Tanzania (2017-242-NA-2017-87).

## DATA ACCESSIBILITY

Should the manuscript be accepted, all data used in this study along with R codes will be made available and archived in the public repository Dryad, the DOI will be included at the end of the article.

## ACKNOWLEDGMENTS

This work was financed by the FNS and ANR Project ‘REEFISH’ no. 310030E-164294.

We thank the local authorities for issuing permits to collect samples and helping with field logistics in the Maldives, Mayotte (Comoros), Seychelles and Mafia Island (Tanzania). We are very grateful to the local staff members, namely Amin Abdallah (Mafia Island Marine Park) and Clara Belmont, Maria Cedras and Rodney Melanie (Seychelles Fishing Authority). We also thank the crew of L’Amitié (Seychelles Fishing Authority) and Ahmed Evaan from Hope Cruiser (Maldives), as well as the Mafia Island Diving center, in particular David von Helldorff, Danielle Keates and Hamis Mjoc, for support and assistance in the field. We are grateful to all the local students who joined and helped during our expeditions, namely Luluesther Samwel, Lucas Simon, Eliad Lukuwi, Oscar Joseph Ngido, Peter Majengo, Ipyana Adamssony and Stephanie Marie. We additionally thank Séverine Albouy, Patrice Descombes, Anna Marcionetti, Joris Bertrand, Nadine Sandau, Caroline Cosnard and Loïc Chalmandrier for help in various field missions and Claudia Michel, Silvia Kobel, Aria Minder, Camille Pitteloud and Livia Gerber for guidance during preparation of the libraries. All the genetic data produced and analyzed in this paper have no commercial value and were generated in collaboration with the Genetic Diversity Centre (GDC), ETH Zurich.

