## Supplementary material for "Species ecology explains the various spatial components of genetic diversity in tropical reef fishes": Biorxiv_Donati_et_al_SuppMat.pdf

**This supporting information contains:**

**List of Tables**

**table S1.** List of studies testing the association between ecological traits and genetic diversity.

**References for table S1**

**table S2.** Taxonomic, sampling and ecological trait information of the 20 genotyped tropical reef fish species.

**table S3.** Phylogenetic signal investigated for functional traits and $\overline{\alpha}$, $\beta$ and $\gamma$ components of genetic diversity.

**table S4.** Genotyping parameter selection used in the *de novo* assembly.

**table S5.** SNP information.

**table S6.** Values of multiplicative and additive partitioning frameworks of genetic diversity.

**table S7.** Genetic α diversity calculated according to the multiplicative framework for biodiversity partitioning.

**table S8.** Genetic $\beta$ diversity estimated between populations as pairwise $F_{ST}$ according to Nei (1987).

**table S9.** Ordinary least squares (OLS) model results for the $\overline{\alpha}$, β and γ components of genetic diversity and the ecological traits investigated.

**table S10**: Results of the Ordinary Least Squares (OLS) and phylogenetic least square (PGLS) models relating the $\overline{\alpha}$, $\beta$ and $\gamma$ components of genetic diversity to the PCA axes,

**List of Figures**

**figure S1.** Pearson correlation matrix between continuous traits and genetic $\overline{\alpha}$, $\beta$ and $\gamma$ diversity for both multiplicative and additive partitioning frameworks of biodiversity.

**figure S2.** Pearson correlation matrix between all metrics of genetic $\beta$ diversity.

**figure S3.** Principal Component Analysis (PCA) plots on genome-wide SNP data for the 20 tropical reef fishes of the Western Indian Ocean considered in our analyses.

**figure S4.** Spatial variation in genetic $\alpha$ diversity across the four sites and across species (*a*) and genetic $\beta$ diversity across the sites in relation to spatial distance (*b*).

**figure S5.** Results of the Principal Component Analysis (PCA) performed on the trait data set for the studied 20 species.

**Supporting methods**

**References**

### Tables

**table S1.** List of studies published since 2009 in which the association between ecological traits and genetic diversity was tested for a minimum of 10 species. The objective of all these studies was to detect the influence of ecological traits and in some cases also of the geography and the environment on $\alpha$, $\beta$ or $\gamma$ levels of genetic diversity. The list was established from a bibliographic search on ISI WEB (September 2020) using the following combination of keywords: "genetic diversity and biological trait and multi-species"; "genetic diversity and ecological trait and multi-species"; "genetic diversity and life history and multi-species"; "genetic structure and ecological trait and multi-species"; "genetic structure and life history and multi-species".

| Organisms | Ecosystem | Study area | n | Markers | Genetic diversity | Conclusion | Ref |
|---|---|---|---|---|---|---|---|
| Animals | All | Global | 76 | nuc: SNP | $\gamma$ | Strong correlation between genetic diversity and life-history traits (body mass, longevity, reproduction). | [1] |
| Animals | Marine, coral reefs | Hawaii | 35 | mtDNA, nuc: msat; intron | $\beta$ | Dispersal ability, taxonomy (i.e. fish vs. invertebrate) and habitat specialization mainly influence genetic differentiation. | [2] |
| Animals | Marine, coral reefs | Hawaii | 47 | mtDNA, nuc: msat | $\alpha$, $\beta$ | Genetic diversity is influenced by depth range and herbivory. | [3] |
| Fish | Freshwater | Global | 12 | msat | $\alpha$ | Meta-analysis: increase in intraspecific genetic diversity is influenced by downstream-biased dispersal, colonization processes and habitat availability. | [4] |
| Fish | Marine | Mediterranean Sea | 31 | mtDNA, nuc: msat | $\alpha$ | Meta-analysis: vertical distribution, migration type and average body length influence genetic diversity. | [5] |
| Fish | Freshwater | Portugal | 17 | mtDNA | $\alpha$, $\beta$ | Contemporary determinants of species' intrinsic traits and landscape features play a more significant role than historical factors in genetic diversity. | [6] |

| | | | | | | | |
|---|---|---|---|---|---|---|---|
| Fish | Marine – Freshwater | Global | 463 | nuc: msat | *α* | Meta-analysis: age maturity and fecundity are negatively related to genetic variation / differences between habitats (marine vs. freshwater). | [7] |
| Fish | Marine, coral reefs | Moorea | 13 | nuc: SNP | *γ* | Trophic ecology influences genetic diversity. | [8] |
| Fish | Marine, coral reefs | Global | 91 | mtDNA, nuc: msat; allozyme | *β* | The reproductive strategy (benthic guarded or pelagic spawned) of reef fishes is a strong predictor of population genetic structure. | [9] |
| Fish | Marine, coral reefs | Pacific Ocean | 11 | mtDNA | *α, β* | Life-history traits are a poor predictor of population structure and migration rates. Isolation and habitat area are the most influential drivers of genetic diversity. | [10] |
| Cetacean | Marine | Global | 42 | mtDNA, nuc: msat | *γ* | Nuclear diversity is correlated with population size, while mitochondrial diversity is influenced by the range and social structure of the species. | [11] |
| Butterfly | Terrestrial | Europe | 38 | RNAseq | *γ* | Genetic diversity correlates negatively with body size and positively with the length of the genetic map. | [12] |
| Butterfly | Terrestrial | Western Central Europe | 307 | mtDNA | *α* | Spatial differentiation is negatively correlated with traits determining dispersibility. | [13] |
| Birds | Terrestrial | Amazonia | 20 | nuc: SNP [a] | *α* | Genetic diversity is associated with habitat type. | [14] |
| Birds | Terrestrial | Amazonia | 40 | mtDNA (CytB) | *β* | Genetic diversity is associated with forest stratum, which is expected to reflect differences in dispersal ability between bird species. | [15] |

| | | | | | | | |
|---|---|---|---|---|---|---|---|
| Birds | Terrestrial | Global | 72 | nuc: msat | $\alpha$ | Genetic diversity is associated with habitat type (terrestrial vs. aquatic) and body mass. | [16] |
| Plants | Terrestrial | Australia | 118 | nuc: msat; Allozyme | $\alpha, \beta$ | Genetic diversity is associated with range size, growth form, abundance and biome. Genetic differentiation is associated with the abundance and distribution of populations across the range | [17] |
| Amphibians | Terrestrial | Canada, US | 299 | mtDNA | $\alpha$ | Genetic diversity is associated with phylogenetic diversity, sample size and some climatic predictors, but no effect of life history traits (body size, development, breeding habitat, and neoteny). Genetic diversity is greater at lower latitude for salamander | [18] |
| Mammals | Marine and Terrestrial | Global | 95 | Nuc: msat | $\gamma$ | Genetic diversity is correlated with range size in threatened species, and ne effect of habitat, trophic class or body size was detected | [19] |

**table S2.** Taxonomic information, species code (Abrev.), total sampling success (n), ecological traits and abundance data of the 20 genotyped tropical reef fish species, namely adult body size (BS), pelagic larval duration (PLD), adult home range mobility behavior, reproductive guild (Repro), schooling (School., number of individuals per school) and local abundance (Abd.).

| Family | Taxon | Abrev. | n | BS (cm) | PLD (days) | Home range | Repro. | School. | Abd. |
|---|---|---|---|---|---|---|---|---|---|
| Tetraodontidae | *Canthigaster valentini* | *Can_v* | 44 | 11 | 18 | narrow | guarders | < 20 | 266 |
| Carangidae | *Caranx melampygus* | *Car_m* | 15 | 117 | 57.6 | wide | non-guarders | < 20 | 24 |
| Chaetodontidae | *Chaetodon trifasciatus* | *Cha_t* | 48 | 15 | 43 | narrow | non-guarders | < 20 | 112 |
| Pomacentridae | *Chromis atripectoralis* | *Chr_a* | 34 | 12 | 19 | narrow | guarders | > 20 | 1393 |
| Pomacentridae | *Chromis ternatensis* | *Chr_t* | 47 | 10 | 28.5 | narrow | guarders | > 20 | 1012 |
| Pomacentridae | *Chromis weberi* | *Chr_w* | 45 | 13.5 | 31.2 | narrow | guarders | > 20 | 526 |
| Acanthuridae | *Ctenochaetus striatus* | *Cte_s* | 69 | 26 | 43.9 | narrow | non-guarders | > 20 | 446 |
| Pomacentridae | *Dascyllus aruanus* | *Das_a* | 38 | 10 | 21.1 | narrow | guarders | > 20 | 68 |
| Pomacentridae | *Dascyllus carneus* | *Das_c* | 54 | 7 | 24.3 | narrow | guarders | > 20 | 2119 |
| Pomacentridae | *Dascyllus trimaculatus* | *Das_t* | 38 | 11 | 28 | narrow | guarders | < 20 | 1072 |
| Labridae | *Gomphosus caeruleus* | *Gom_c* | 43 | 30 | 56.6 | wide | non-guarders | < 20 | 213 |
| Labridae | *Halichoeres hortulanus* | *Hal_h* | 43 | 27 | 32.5 | wide | non-guarders | < 20 | 207 |
| Labridae | *Hemigymnus fasciatus* | *Hem_f* | 42 | 30 | 25.8 | wide | non-guarders | < 20 | 39 |
| Lutjanidae | *Lutjanus kasmira* | *Lut_k* | 33 | 40 | 31 | wide | non-guarders | > 20 | 28 |
| Holocentridae | *Myripristis violacea* | *Myr_v* | 38 | 35 | 60.4 | wide | non-guarders | < 20 | 69 |
| Acanthuridae | *Naso brevirostris* | *Nas_b* | 20 | 60 | 79.7 | wide | non-guarders | > 20 | 11 |
| Monacanthidae | *Oxymonacanthus longirostris* | *Oxy_l* | 57 | 12 | 26 | narrow | guarders | < 20 | 125 |
| Mullidae | *Parupeneus macronemus* | *Par_m* | 55 | 40 | 41.8 | wide | non-guarders | < 20 | 101 |
| Serranidae | *Pseudanthias squamipinnis* | *Pse_s* | 47 | 15 | 26 | narrow | non-guarders | > 20 | 4422 |
| Zanclidae | *Zanclus cornutus* | *Zan_c* | 42 | 23 | 57.9 | wide | non-guarders | < 20 | 69 |

**table S3.** Phylogenetic signal investigated for ecological traits and $\alpha$, $\beta$ and $\gamma$ components of genetic diversity with both additive ($H_S$, $H_T$, $D_{ST}$) and multiplicative ($J_S$, $J_T$, $J_{ST}$) partitioning frameworks of biodiversity. We applied the $\lambda$ metric for continuous traits [20] and the D-statistic [21] for categorical traits.

| Variable | Metric of phylogenetic signal | Value |
|---|---|---|
| Abundance | $\lambda$ | 6.616e-05 |
| Body size | $\lambda$ | 0.999 |
| PLD | $\lambda$ | 0.999 |
| Home range | D-statistic | -0.294 |
| Reproduction | D-statistic | -2.182 |
| Schooling | D-statistic | 0.065 |
| $H_S$ | $\lambda$ | 0.999 |
| $H_T$ | $\lambda$ | 0.999 |
| $D_{ST}$ | $\lambda$ | 6.616e-05 |
| $J_S$ | $\lambda$ | 0.999 |
| $J_T$ | $\lambda$ | 0.999 |
| $J_{ST}$ | $\lambda$ | 6.616e-05 |

**table S4.** Parameter selection (coverage, shared loci and percentage of sequence identity) used in the *de novo* assembly.

| Taxon | De novo assembly parameters: [coverage - shared loci - sequence identity] | Restriction: site check / no site check |
|---|---|---|
| *Chaetodon trifasciatus* | 3-4-90 | site check |
| *Canthigaster valentini* | 3-4-95 | site check |
| *Dascyllus aruanus* | 3-4-95 | site check |
| *Dascyllus carneus* | 3-3-92 | site check |
| *Dascyllus trimaculatus* | 3-4-95 | site check |
| *Pseudanthias squamipinnis* | 5-2-95 | site check |
| *Zanclus cornutus* | 3-4-95 | site check |
| *Chromis ternatensis* | 3-3-92 | site check |
| *Chromis weberi* | 3-3-95 | site check |
| *Chromis atripectoralis* | 3-3-92 | site check |
| *Gomphosus caeruleus* | 2-2-90 | no check |
| *Halichoeres hortulanus* | 2-2-90 | site check |
| *Hemigymnus fasciatus* | 3-2-90 | no check |
| *Lutjanus kasmira* | 3-2-90 | no check |
| *Myripristis violacea* | 3-3-95 | no check |
| *Parupeneus macronema* | 3-3-92 | no check |
| *Oxymonacanthus longirostris* | 2-2-90 | site check |
| *Naso brevirostris* | 3-4-95 | site check |
| *Caranx melampygus* | 3-4-90 | site check |
| *Ctenochaetus striatus* | 3-4-90 | site check |

**table S5.** Sample size and SNP data on 20 tropical reef fish species in the Western Indian Ocean. (Left) Total sampling refers to the complete data set; (Right) standardized re-sampling refers to the standardized and resampled (x999) data set, where the maximum *n* per sample site was 10 (i.e. the median of the complete data set). Sampling location codes: Mafia Island (MF), Maldives (MV), Mayotte Island (MY) and Seychelles (SC). The number of total, filtered and resampled SNPs are also indicated.

| Taxon | Total sampling | | | | | | Standardized re-sampling | | | | |
|---|---|---|---|---|---|---|---|---|---|---|---|
| | MF | MV | MY | SC | Total SNPs | Filtered SNPs | MF | MV | MY | SC | Resampled SNPs |
| *Canthigaster valentini* | 10 | 10 | 14 | 10 | 339809 | 15080 | 10 | 10 | 10 | 10 | 4479 |
| *Caranx melampygus* | 3 | 4 | 6 | 2 | 107395 | 15431 | 3 | 4 | 6 | 2 | 4479 |
| *Chaetodon trifasciatus* | 11 | 12 | 15 | 10 | 327595 | 20950 | 10 | 10 | 10 | 10 | 4479 |
| *Chromis atripectoralis* | 3 | 9 | 9 | 13 | 1255718 | 7225 | 3 | 9 | 9 | 10 | 4479 |
| *Chromis ternatensis* | 10 | 14 | 14 | 9 | 873075 | 17369 | 10 | 10 | 10 | 9 | 4479 |
| *Chromis weberi* | 10 | 10 | 13 | 12 | 926904 | 14370 | 10 | 10 | 10 | 10 | 4479 |
| *Ctenochaetus striatus* | 10 | 15 | 40 | 4 | 697607 | 9664 | 10 | 10 | 10 | 4 | 4479 |
| *Dascyllus aruanus* | 11 | 12 | 7 | 8 | 986800 | 22105 | 10 | 10 | 7 | 8 | 4479 |
| *Dascyllus carneus* | 8 | 17 | 20 | 9 | 1086702 | 16260 | 8 | 10 | 10 | 9 | 4479 |
| *Dascyllus trimaculatus* | 8 | 12 | 8 | 10 | 682522 | 38934 | 8 | 10 | 8 | 10 | 4479 |
| *Gomphosus caeruleus* | 9 | 12 | 12 | 10 | 854088 | 6253 | 9 | 10 | 10 | 10 | 4479 |
| *Halichoeres hortulanus* | 7 | 14 | 13 | 9 | 242929 | 18934 | 7 | 10 | 10 | 9 | 4479 |
| *Hemigymnus fasciatus* | 10 | 11 | 11 | 10 | 229050 | 15606 | 10 | 10 | 10 | 10 | 4479 |
| *Lutjanus kasmira* | 9 | 9 | 11 | 4 | 806613 | 7420 | 9 | 9 | 10 | 4 | 4479 |
| *Myripristis violacea* | 10 | 9 | 7 | 12 | 463771 | 14890 | 10 | 9 | 7 | 10 | 4479 |
| *Naso brevirostris* | 12 | 1 | 3 | 4 | 357378 | 15134 | 10 | 1 | 3 | 4 | 4479 |
| *Oxymonacanthus longirostris* | 8 | 11 | 35 | 3 | 111506 | 9679 | 8 | 10 | 10 | 3 | 4479 |
| *Parupeneus macronema* | 9 | 17 | 22 | 7 | 635207 | 7432 | 9 | 10 | 10 | 7 | 4479 |
| *Pseudanthias squamipinnis* | 13 | 15 | 8 | 11 | 696705 | 4479 | 10 | 10 | 8 | 10 | 4479 |
| *Zanclus cornutus* | 9 | 12 | 11 | 10 | 274244 | 17590 | 9 | 10 | 10 | 10 | 4479 |

**table S6. Values of multiplicative and additive partitioning frameworks of genetic diversity.** To quantify the different $\overline{\alpha}$, $\beta$ and $\gamma$ components we used the partitioning for true diversities proposed by Jost [22] and applied to genetic data, expressed as follows: $J_T = J_S \times J_{ST}$ where $J_S$ represents the within-population genetic component ($\overline{\alpha}$), $J_{ST}$ represents the between-population ($\beta$) component and $J_T$ the overall genetic diversity ($\gamma$). We compared the multiplicative framework with an additive one expressed as $\beta = \gamma - \overline{\alpha}$, for which we used the mean heterozygosity ($H_S$) as a measure of $\overline{\alpha}$ diversity and overall gene diversity ($H_T$) as a measure of $\gamma$ diversity. The $\beta$ diversity is the equivalent of the $D_{ST}$ metric, where $D_{ST} = H_T - H_S$ [23]. All of these metrics were calculated for each species and each of the 999 x 4479 SNP data sets.

| | **Multiplicative partitioning framework** | | | **Additive partitioning framework** | | |
|---|---|---|---|---|---|---|
| **Taxon** | $J_T$ ($\gamma$) | $J_S$ ($\overline{\alpha}$) | $J_{ST}$ ($\beta$) | $H_T$ ($\gamma$) | $H_S$ ($\overline{\alpha}$) | $D_{ST}$ ($\beta$) |
| *Canthigaster valentini* | 1.3785 | 1.3743 | 1.0030 | 0.2746 | 0.2723 | 0.0022 |
| *Caranx melampygus* | 1.4721 | 1.4739 | 1.0000 | 0.3207 | 0.3207 | 0.0000 |
| *Chaetodon trifasciatus* | 1.3755 | 1.3735 | 1.0015 | 0.2730 | 0.2719 | 0.0011 |
| *Chromis atripectoralis* | 1.3702 | 1.3655 | 1.0034 | 0.2702 | 0.2677 | 0.0025 |
| *Chromis ternatensis* | 1.3520 | 1.3459 | 1.0045 | 0.2603 | 0.2570 | 0.0033 |
| *Chromis weberi* | 1.3375 | 1.3317 | 1.0043 | 0.2523 | 0.2491 | 0.0032 |
| *Ctenochaetus striatus* | 1.3720 | 1.3717 | 1.0002 | 0.2711 | 0.2710 | 0.0002 |
| *Dascyllus aruanus* | 1.3741 | 1.3654 | 1.0064 | 0.2722 | 0.2676 | 0.0046 |
| *Dascyllus carneus* | 1.3482 | 1.3466 | 1.0011 | 0.2582 | 0.2574 | 0.0008 |
| *Dascyllus trimaculatus* | 1.3630 | 1.3615 | 1.0011 | 0.2663 | 0.2655 | 0.0008 |
| *Gomphosus caeruleus* | 1.3722 | 1.3716 | 1.0004 | 0.2713 | 0.2709 | 0.0003 |
| *Halichoeres hortulanus* | 1.4112 | 1.4097 | 1.0011 | 0.2914 | 0.2906 | 0.0008 |
| *Hemigymnus fasciatus* | 1.4441 | 1.4241 | 1.0140 | 0.3075 | 0.2978 | 0.0097 |
| *Lutjanus kasmira* | 1.4135 | 1.4131 | 1.0003 | 0.2925 | 0.2923 | 0.0002 |
| *Myripristis violacea* | 1.3785 | 1.3783 | 1.0001 | 0.2746 | 0.2745 | 0.0001 |
| *Naso brevirostris* | 1.4466 | 1.4457 | 1.0006 | 0.3087 | 0.3083 | 0.0004 |
| *Oxymonacanthus longirostris* | 1.3957 | 1.3782 | 1.0127 | 0.2835 | 0.2744 | 0.0091 |
| *Parupeneus macronemus* | 1.3686 | 1.3625 | 1.0045 | 0.2693 | 0.2660 | 0.0033 |
| *Pseudanthias squamipinnis* | 1.3134 | 1.3099 | 1.0026 | 0.2386 | 0.2366 | 0.0019 |
| *Zanclus cornutus* | 1.3785 | 1.3783 | 1.0001 | 0.2746 | 0.2745 | 0.0001 |

**table S7.** Genetic $\alpha$ diversity calculated according to the multiplicative framework for biodiversity partitioning $J_S$= $1/(1 - H_S)$ where $H_S$ is the expected heterozygosity. $J_S$ was calculated for a standardized number of individuals per population over 999 randomizations.

| **Genetic $\alpha$ diversity** | **$J_S$ Maldives** | | **$J_S$ Seychelles** | | **$J_S$ Mafia Island** | | **$J_S$ Mayotte** | |
|---|---|---|---|---|---|---|---|---|
| **Taxon** | mean | sd | mean | sd | mean | sd | mean | sd |
| *Canthigaster valentini* | 1.366 | 0.004 | 1.380 | 0.004 | 1.378 | 0.004 | 1.374 | 0.004 |
| *Caranx melampygus* | 1.474 | 0.005 | 1.471 | 0.006 | 1.482 | 0.004 | 1.466 | 0.008 |
| *Chaetodon trifasciatus* | 1.370 | 0.004 | 1.376 | 0.004 | 1.374 | 0.004 | 1.373 | 0.004 |
| *Chromis atripectoralis* | 1.339 | 0.002 | 1.378 | 0.002 | 1.361 | 0.003 | 1.384 | 0.002 |
| *Chromis ternatensis* | 1.326 | 0.004 | 1.352 | 0.004 | 1.354 | 0.004 | 1.353 | 0.003 |
| *Chromis weberi* | 1.312 | 0.004 | 1.338 | 0.003 | 1.338 | 0.004 | 1.338 | 0.003 |
| *Ctenochaetus striatus* | 1.368 | 0.003 | 1.374 | 0.004 | 1.371 | 0.003 | 1.375 | 0.003 |
| *Dascyllus aruanus* | 1.338 | 0.004 | 1.374 | 0.004 | 1.376 | 0.004 | 1.374 | 0.005 |
| *Dascyllus carneus* | 1.342 | 0.003 | 1.346 | 0.004 | 1.349 | 0.004 | 1.348 | 0.003 |
| *Dascyllus trimaculatus* | 1.357 | 0.004 | 1.365 | 0.004 | 1.365 | 0.005 | 1.359 | 0.005 |
| *Gomphosus caeruleus* | 1.371 | 0.002 | 1.376 | 0.002 | 1.367 | 0.002 | 1.372 | 0.002 |
| *Halichoeres hortulanus* | 1.406 | 0.004 | 1.412 | 0.004 | 1.409 | 0.005 | 1.412 | 0.004 |
| *Hemigymnus fasciatus* | 1.384 | 0.004 | 1.436 | 0.004 | 1.441 | 0.004 | 1.437 | 0.004 |
| *Lutjanus kasmira* | 1.416 | 0.002 | 1.412 | 0.003 | 1.412 | 0.002 | 1.413 | 0.002 |
| *Myripristis violacea* | 1.376 | 0.004 | 1.377 | 0.004 | 1.378 | 0.004 | 1.382 | 0.004 |
| *Naso brevirostris* | NA | NA | 1.451 | 0.005 | 1.443 | 0.004 | 1.448 | 0.006 |
| *Oxymonacanthus longirostris* | 1.353 | 0.003 | 1.394 | 0.004 | 1.397 | 0.003 | 1.370 | 0.005 |
| *Parupeneus macronemus* | 1.362 | 0.003 | 1.360 | 0.003 | 1.364 | 0.003 | 1.364 | 0.003 |
| *Pseudanthias squamipinnis* | 1.315 | 0 | 1.310 | 0 | 1.309 | 0 | 1.305 | 0 |
| *Zanclus cornutus* | 1.378 | 0.004 | 1.380 | 0.004 | 1.379 | 0.004 | 1.377 | 0.004 |

**table S8.** Genetic $\beta$ diversity estimated between populations as pairwise $F_{ST}$ according to Nei [23]. $F_{ST}$ values were calculated for a standardized number of individuals per population over 999 x 4479 SNP randomizations. Populations are labeled as: Maldives (MV), Mafia Island (MF), Mayotte Island (MY) and Seychelles (SC).

| **Genetic β diversity** | **MV_MF** | | **MV_MY** | | **MV_SC** | | **MF_SC** | | **MY_SC** | | **MF_MY** | |
|---|---|---|---|---|---|---|---|---|---|---|---|---|
| **Genus species** | **mean** | **sd** | **mean** | **sd** | **mean** | **sd** | **mean** | **sd** | **mean** | **sd** | **mean** | **sd** |
| *Canthigaster valentini* | 0.021 | 0.002 | 0.021 | 0.002 | 0.022 | 0.002 | 0.001 | 0.001 | 0.001 | 0.001 | 0 | 0 |
| *Caranx melampygus* | 0 | 0 | 0 | 0 | 0 | 0 | 0 | 0 | 0 | 0 | 0 | 0 |
| *Chaetodon trifasciatus* | 0.008 | 0.002 | 0.01 | 0.002 | 0.01 | 0.002 | 0.002 | 0.001 | 0.002 | 0.001 | 0.001 | 0.001 |
| *Chromis atripectoralis* | 0.028 | 0.002 | 0.018 | 0.001 | 0.017 | 0.001 | 0.009 | 0.001 | 0.003 | 0.001 | 0.002 | 0.001 |
| *Chromis ternatensis* | 0.035 | 0.002 | 0.037 | 0.002 | 0.036 | 0.002 | 0 | 0 | 0 | 0 | 0 | 0 |
| *Chromis weberi* | 0.034 | 0.002 | 0.034 | 0.002 | 0.034 | 0.002 | 0.001 | 0.001 | 0 | 0 | 0 | 0 |
| *Ctenochaetus striatus* | 0.006 | 0.001 | 0.003 | 0 | 0 | 0 | 0 | 0 | 0 | 0 | 0 | 0 |
| *Dascyllus aruanus* | 0.042 | 0.002 | 0.042 | 0.002 | 0.047 | 0.002 | 0.001 | 0.001 | 0.001 | 0.002 | 0.001 | 0.002 |
| *Dascyllus carneus* | 0.006 | 0.002 | 0.010 | 0.001 | 0.01 | 0.001 | 0 | 0 | 0.001 | 0.001 | 0 | 0 |
| *Dascyllus trimaculatus* | 0.008 | 0.002 | 0.007 | 0.002 | 0.006 | 0.001 | 0.002 | 0.001 | 0 | 0 | 0.001 | 0.001 |
| *Gomphosus caeruleus* | 0.003 | 0.001 | 0 | 0.001 | 0.002 | 0.001 | 0.002 | 0.001 | 0.001 | 0.001 | 0.003 | 0.001 |
| *Halichoeres hortulanus* | 0.006 | 0.001 | 0.008 | 0.001 | 0.007 | 0.001 | 0.001 | 0.001 | 0 | 0.001 | 0 | 0 |
| *Hemigymnus fasciatus* | 0.078 | 0.002 | 0.08 | 0.002 | 0.079 | 0.002 | 0.003 | 0.001 | 0.004 | 0.001 | 0.001 | 0.001 |
| *Lutjanus kasmira* | 0 | 0 | 0 | 0 | 0.003 | 0.001 | 0.004 | 0.001 | 0.001 | 0.001 | 0.001 | 0.001 |
| *Myripristis violacea* | 0.002 | 0.001 | 0 | 0 | 0 | 0 | 0.001 | 0.001 | 0 | 0 | 0 | 0 |
| *Naso brevirostris* | 0.001 | 0.007 | 0.004 | 0.01 | 0.003 | 0.008 | 0.001 | 0.002 | 0 | 0 | 0.003 | 0.002 |
| *Oxymonacanthus longirostris* | 0.056 | 0.001 | 0.066 | 0.003 | 0.060 | 0.002 | 0.005 | 0.001 | 0.033 | 0.002 | 0.033 | 0.002 |
| *Parupeneus macronemus* | 0.028 | 0.001 | 0.034 | 0.001 | 0.037 | 0.002 | 0 | 0 | 0 | 0 | 0 | 0 |
| *Pseudanthias squamipinnis* | 0.020 | 0 | 0.021 | 0 | 0.021 | 0 | 0.002 | 0 | 0 | 0 | 0 | 0 |
| *Zanclus cornutus* | 0 | 0 | 0.002 | 0.001 | 0 | 0 | 0 | 0 | 0.001 | 0.001 | 0.001 | 0.001 |

**table S9.** Results of the ordinary least squares (OLS) models for the $\overline{\alpha}$, $\beta$ and $\gamma$ components of genetic diversity and the six ecological traits investigated. For quantitative traits, we report the significance of the PGLS regression coefficient (slope), while for qualitative traits we report the significance of the Fisher statistic (F) derived from an ANOVA applied to the OLS model. Significant variables ($P$ < 0.05) are given in bold.

| **$\overline{\alpha}$ genetic diversity** | OLS | | | | | |
|---|---|---|---|---|---|---|
| | $R^2$ | Coef. | F | *P* | AICc | $W_{AICc}$ |
| **Body size** | **0.594** | **0.0416** | **-** | **0.00007** | **-88.345** | **0.143** |
| Pelagic larval duration | 0.243 | 0.0111 | - | 0.0273 | -75.866 | 0 |
| **Abundance** | **0.661** | **-0.0191** | **-** | **0.00001** | **-91.909** | **0.851** |
| **Home range** | **0.4393** | **-** | **14.100** | **0.00145** | **-81.872** | **0.006** |
| Schooling | 0.0975 | - | 1.945 | 0.1801 | -72.354 | 0 |
| Reproduction | 0.201 | - | 4.515 | 0.0477 | -74.779 | 0 |
| **β genetic diversity** | OLS | | | | | |
| | $R^2$ | Coef. | F | *P* | AICc | $W_{AICc}$ |
| Body size | 0.0774 | -0.00156 | - | 0.235 | -162.327 | 0.103 |
| **Pelagic larval duration** | **0.228** | **-0.00012** | **-** | **0.0335** | **-165.879** | **0.608** |
| Abundance | 0.0004 | -0.00016 | - | 0.779 | -160.805 | 0.048 |
| Home range | 0.0309 | - | 0.574 | 0.4585 | -161.343 | 0.063 |
| Schooling | 0.0135 | - | 0.248 | 0.625 | -160.989 | 0.053 |
| Reproduction | 0.0959 | - | 1.910 | 0.184 | -162.733 | 0.126 |
| **γ genetic diversity** | OLS | | | | | |
| | $R^2$ | Coef. | F | *P* | AICc | $W_{AICc}$ |
| **Body size** | **0.529** | **0.0395** | **-** | **0.00028** | **-85.069** | **0.027** |
| Pelagic larval duration | 0.178 | 0.00095 | - | 0.0643 | -73.935 | 0 |
| **Abundance** | **0.670** | **-0.0194** | **-** | **0.00001** | **-92.204** | **0.970** |
| **Home range** | **0.403** | **-** | **12.158** | **0.00263** | **-80.348** | **0.003** |
| Schooling | 0.108 | - | 2.175 | 0.158 | -72.308 | 0 |
| Reproduction | 0.162 | - | 3.468 | 0.0790 | -73.549 | 0 |

**table S10**: Results of the Ordinary Least Squares (OLS) and phylogenetic least square (PGLS) models relating the $\overline{\alpha}$, $\beta$ and $\gamma$ components of genetic diversity to the PCA axes, which account for the correlations between the species traits (see Fig. S3). Specifically, the PCA axis 1 represents a gradient of species with varying abundance, body size, home range mobility, reproduction type and PLD. Significant variables ($P < 0.05$) are given in bold. Results revealed a significant and negative association between the $\overline{\alpha}$, and $\gamma$ components of genetic diversity and the PCA axis 1, which confirms that large-bodied species with low regional abundance display the greatest levels of $\overline{\alpha}$ and $\gamma$ genetic diversity.

| **$\overline{\alpha}$ genetic diversity** | OLS | | | PGLS | | |
|---|---|---|---|---|---|---|
| | $R^2$ | Coef. | P | $R^2$ | Coef. | P |
| **PCA axis 1** | **0.4196** | **-5.549** | **0.002** | **0.4132** | **-0.0150** | **0.002** |
| PCA axis 2 | 0.0039 | 0.305 | 0.794 | 0.0073 | 0.0026 | 0.720 |
| ***$\beta$* genetic diversity** | OLS | | | PGLS | | |
| | $R^2$ | Coef. | P | $R^2$ | Coef. | P |
| PCA axis 1 | 0.0685 | 21.490 | 0.265 | 0.0117 | 0.0004 | 0.579 |
| PCA axis 2 | 0.0859 | -13.760 | 0.210 | 0.0073 | 0.0003 | 0.720 |
| **$\gamma$ genetic diversity** | OLS | | | PGLS | | |
| | $R^2$ | Coef. | P | $R^2$ | Coef. | P |
| **PCA axis 1** | **0.3692** | **- 5.169** | **0.004** | **0.3532** | **-0.0144** | **0.005** |
| PCA axis 2 | 0.0003 | 0.0944 | 0.935 | 0.0096 | 0.0031 | 0.680 |

**Figures**

**figure S1.** Pearson correlation matrix between continuous traits for the 20 selected species and genetic $\overline{\alpha}$, $\beta$ and $\gamma$ diversity for both multiplicative ($J_S, J_T, J_{ST}$) and additive partitioning ($H_S, H_T, D_{ST}$) frameworks of biodiversity. Red indicates a significant negative correlation between two variables while blue indicates a significant positive correlation, and the size of the circle represents the significance of the Pearson correlation statistical test. The color gradient (from red to blue) indicates the magnitude of the correlation. White means that the correlation between indicators is not significant according to the Pearson correlation statistical test (PLD: pelagic larval duration).

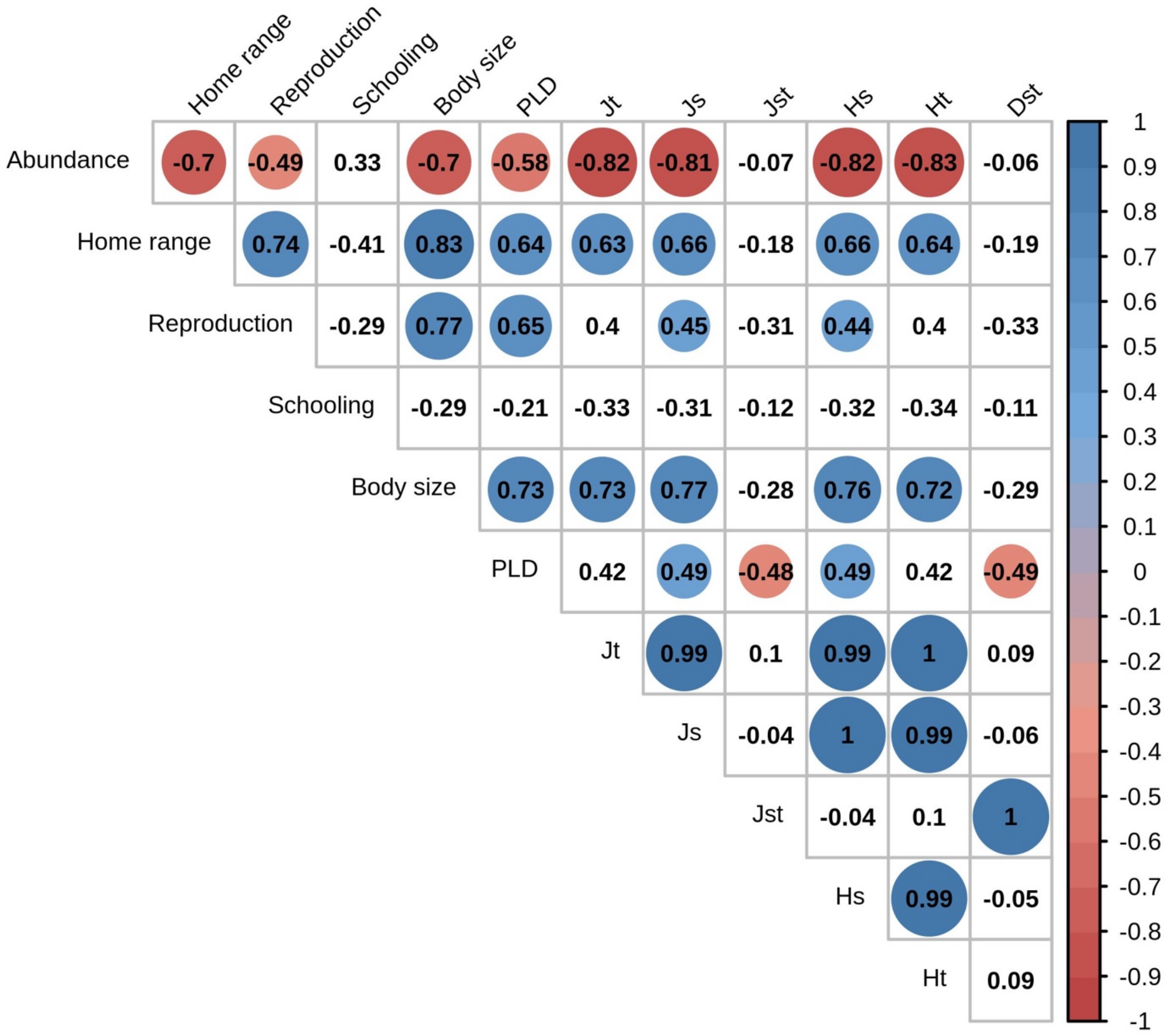

**figure S2.** Pearson correlation matrix between all metrics of $\beta$ diversity. Blue indicates a positive correlation between two variables and the size of the circle represents the significance of the Pearson correlation test. The color gradient (from red to blue) indicates the magnitude of the correlation. Dst' [23], $F_{ST}$ [24], $F_{ST}$', $G_{ST}$, $G_{ST}$', $G_{ST}$'' [25] and *Dest*[22]. Metrics derived from the multiplicative ($J_{ST}$) and additive ($D_{ST}$) partitioning approaches are also presented.

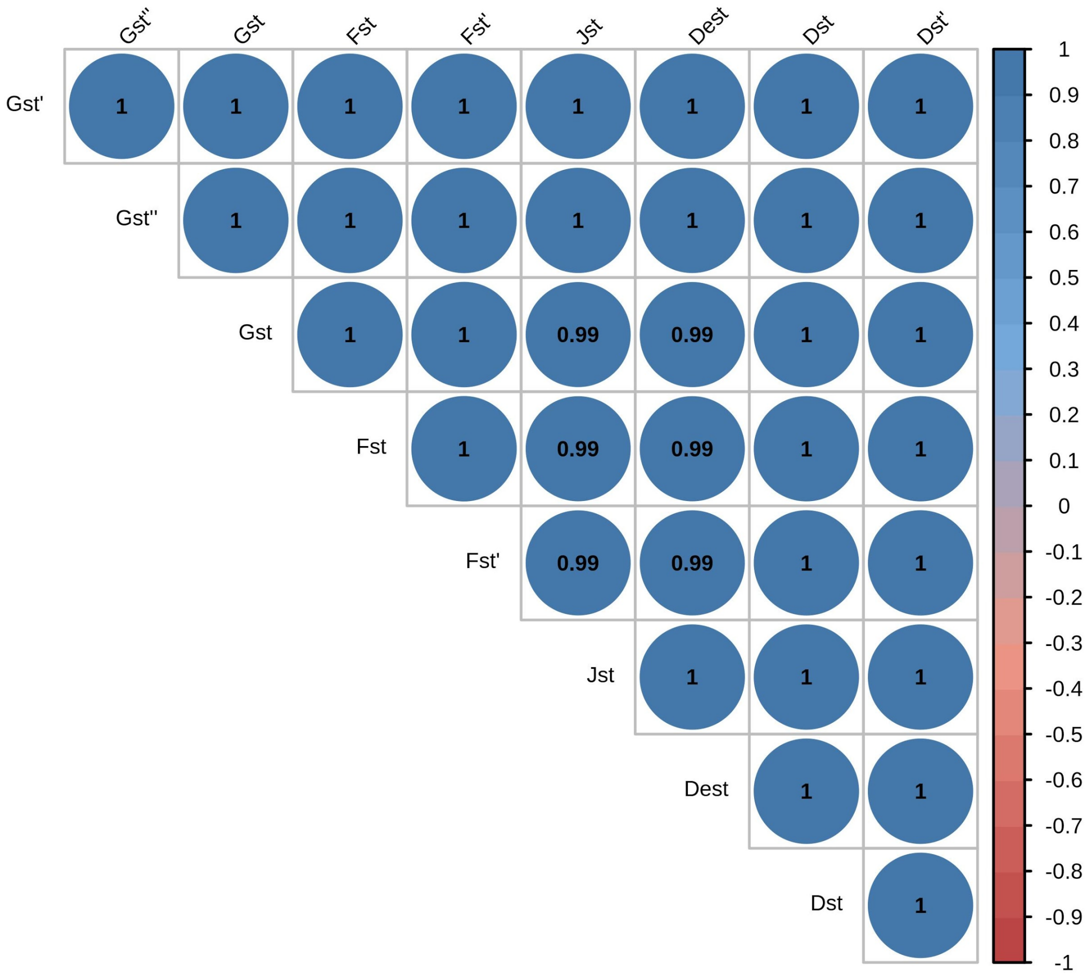

**figure S3.** Results of the Principal Component Analysis (PCA) performed on the trait data set for the 20 studied species. The panel *(a)* represents the correlation circle of traits and the panel *(b)* shows the ordination of species on the plan defined by the two first axes. The first axis reflects a gradient of abundance on the reef, body size, reproductive guild, home range mobility and PLD, with positive coordinates on the PC1 including species with lower abundance, larger body size, higher PLD, higher home range mobility and also adopting a "non-guarder" reproductive strategy. The second axis is only associated with schooling.

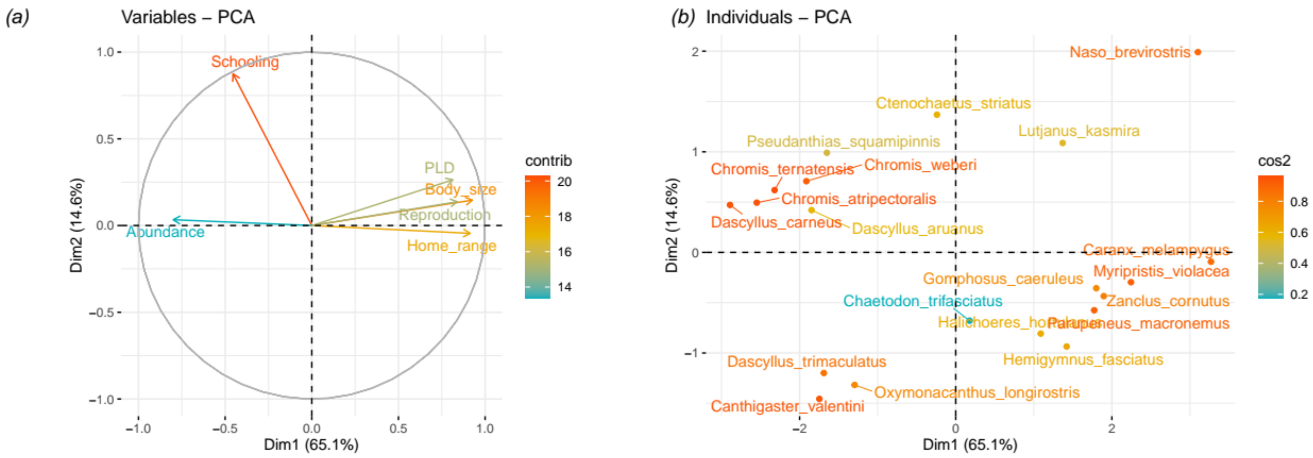

**figure S4.** Spatial variation in $\overline{\alpha}$ diversity across the four sites and across species (a) and $\beta$ diversity across sites in relation to spatial distance (b). Species name abbreviations can be found in table S2. We calculated the pairwise genetic $\beta$ diversity ($F_{ST}$) using the function *genet.dist* in the R package "hierfstat"[26]. To explore the relationship between pairwise dissimilarity and geographical distance, we fitted power-law models with a log-transformed error for each species, which describe the increase in species dissimilarity with spatial distance.

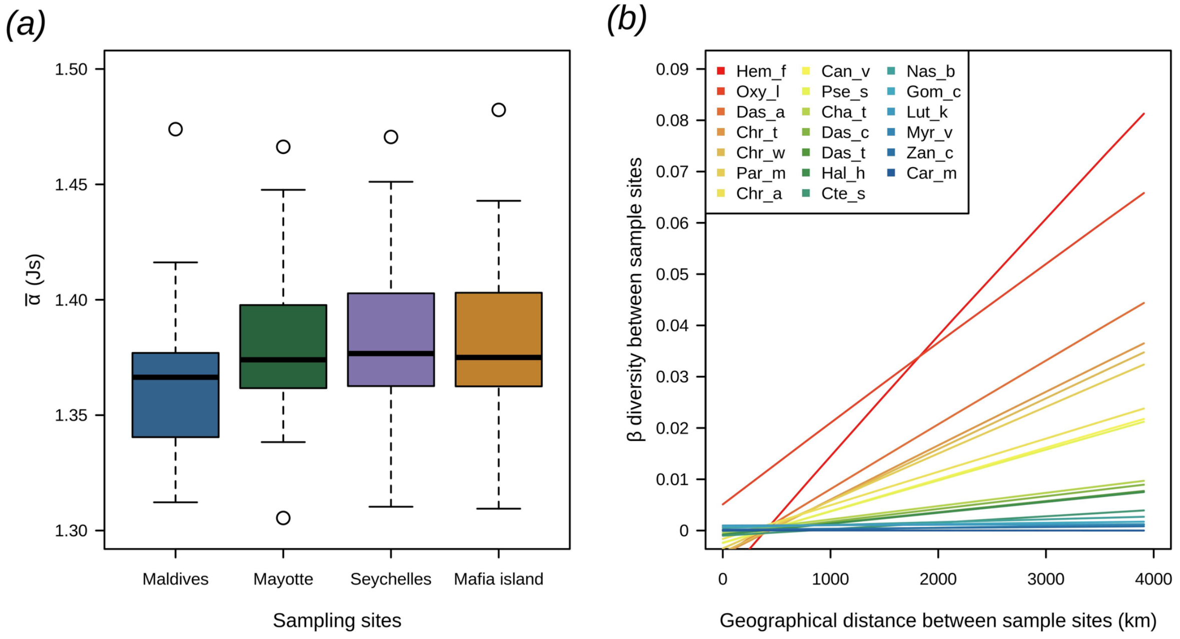

**figure S5.** Principal Component Analysis (PCA) plots on genome-wide SNP data for the 20 tropical reef fishes of the Western Indian Ocean considered in our analyses. The colors of the points and the ellipses represent the different sampling locations, with MV: Maldives, MF: Mafia Island, Sc: Seychelles and MY: Mayotte (MY).

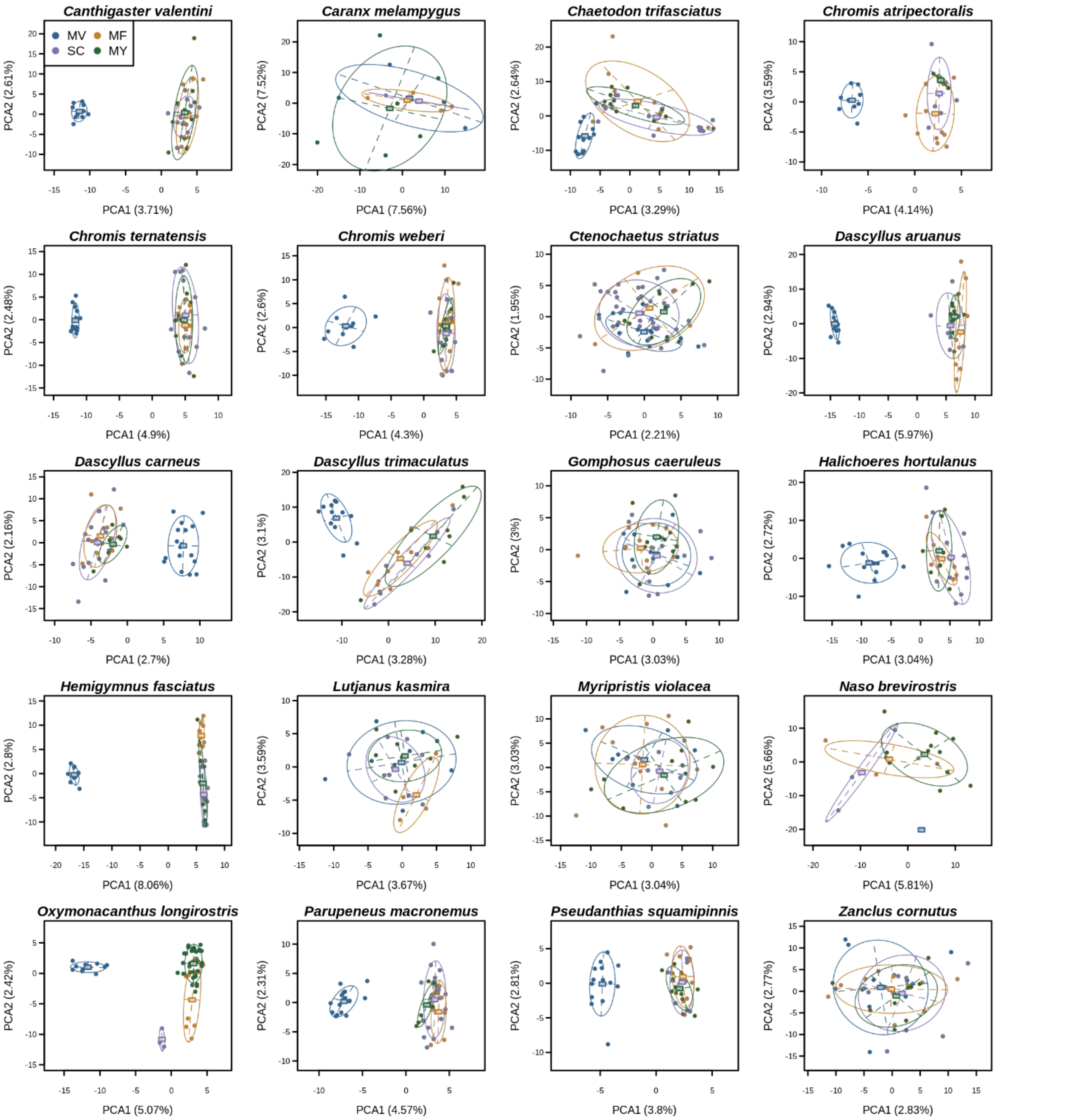

**Supplementary methods**

**Species selection along gradients of ecological traits**

To select 20 tropical reef fish species from the 4 WIO locations, we considered five ecological traits gathered from Fishbase [27] and the literature [8,9,28,29]. These traits include: (i) adult body size, a continuous variable measure (cm) of the total length; (ii) PLD, measured in days; (iii) home range mobility, coded as "narrow" vs. "wide"; (iv) reproductive guild, coded into two simplified categories, "guarders" (which included any type of parental care but mainly species adopting the "benthic-guarding" strategy) vs. "non-guarders" (which excluded any type of parental care and essentially included species adopting an "open water spawning" strategy); and (v) schooling, was coded into two categories for small (< 20 individuals) and large (> 20 individuals) aggregations. These categorical traits were considered as quantitative binary variables (0/1) when running the PCA.
